# Discovery of new senolytics using machine learning

**DOI:** 10.1101/2022.04.26.489505

**Authors:** Vanessa Smer-Barreto, Andrea Quintanilla, Richard J. R. Elliot, John C. Dawson, Jiugeng Sun, Neil O. Carragher, Juan Carlos Acosta, Diego A. Oyarzún

**Affiliations:** Cancer Research UK Edinburgh Centre, MRC Institute of Genetics and Cancer, University of Edinburgh, Crewe Road, Edinburgh, EH4 2XR, UK; Instituto de Biomedicina y Biotecnología de Cantabria (IBBTEC), CSIC-Universidad de Cantabria-SODERCAN. C/ Albert Einstein 22, Santander, 39011, Spain; School of Informatics, University of Edinburgh, 10 Crichton St, Edinburgh, EH8 9AB, UK; School of Biological Sciences, University of Edinburgh, Max Born Crescent, Edinburgh, EH9 3BF, UK; The Alan Turing Institute, 96 Euston Road, London, NW1 2DB, UK

**Keywords:** senescence, machine learning, artificial intelligence, drug discovery

## Abstract

Cellular senescence is a stress response characterised by a permanent cell cycle arrest and a proinflammatory secretome. In addition to its tumour suppressor role, senescence is involved in ageing and promotes many disease processes such as cancer, type 2 diabetes, osteoarthritis, and SARS-CoV-2 infection. There is a growing interest in therapies based on targeted elimination of senescent cells, yet so far only a few such senolytics are known, partly due to the poor grasp of the molecular mechanisms that control the senescence survival programme. Here we report a highly effective machine learning pipeline for the discovery of senolytic compounds. Using solely published data, we trained machine learning algorithms to classify compounds according to their senolytic action. Models were trained on as few as 58 known senolytics against a background of FDA-approved compounds or in late-stage clinical development (2,523 in total). We computationally screened various chemical libraries and singled out top candidates for validation in human lung fibroblasts (IMR90) and lung adenocarcinoma (A549) cell lines. This led to the discovery of three novel senolytics: ginkgetin, oleandrin and periplocin, with potency comparable to current senolytics and a several hundred-fold reduction in experimental screening costs. Our work demonstrates that machine learning can take maximum advantage of existing drug screening data, paving the way for new open science approaches to drug discovery for senescence-associated diseases.

## Introduction

Senescence is a cellular state characterised by permanent cell cycle arrest, macromolecular damage and a range of metabolic alterations [1]. The senescent phenotype can be triggered by different stressors such as replicative exhaustion, chemotherapeutic agents, oncogenic activation, and radiation [2]. Senescent cells affect tissue microenvironment in multiple ways, both beneficial and deleterious [3]. For example, senescence aids mammalian embryonic development, promotes wound healing and stemness [4,5], and is a potent tumour suppression mechanism that restrains the growth of cells in danger of malignant alterations [6]. Conversely, senescent cells also promote tumorigenesis and various age-related malignancies due to the secretion of a complex set of proteins known as the senescence-associated secretory phenotype (SASP) [7,8]. Besides their role in cancer and ageing [9], the senescent programme has been linked to adverse effects in a broad range of conditions, including osteoporosis, osteoarthritis, pulmonary fibrosis, SARS-CoV-2 infection, hepatic steatosis, and neurodegeneration [10]. As a result, there is a growing interest in the discovery of new senolytics, i.e. therapeutic agents that selectively target senescent cells for elimination [10].

Senolytics have shown substantial promise in ameliorating symptoms of many conditions in mice [11–20]. Despite these encouraging results, to date there are few known compounds with proven senolytic action [11–19,21–26], and only two compounds have shown efficacy in clinical trials (dasatinib and quercetin in combination therapy [27]). Some of the most scrutinised senolytics were identified by targeting anti-apoptotic proteins upregulated in senescence, such as the Bcl-2 family inhibitors navitoclax [26] and ABT-737 [16]. Other senolytics were discovered through panel screens [17] and, more recently, high-throughput screens have identified cardiac glycosides such as ouabain [13] and digoxin [15] as potent senolytic agents. A key challenge for senolytic therapies to succeed is that many such compounds display cell-type specific action. In addition, certain senolytics that work well for one cell-type are highly toxic against other non-senescent cell-types [19,22]. In the case of cancer therapies, most senolytics target pathways that are mutated in cancer, limiting their applicability as therapeutic agents [28]. As a result, there is a need for new strategies to identify compounds with senolytic action.

Computational screens have been employed for decades in drug discovery [29–32]. Artificial Intelligence (AI) methods, in particular, have been widely adopted by industrial and academic laboratories due to their ability to detect hidden patterns in large collections of chemical data, which can be used to narrow down the search space for active molecules. These approaches typically rely on a combination of molecular dynamics and sophisticated pipelines to navigate the space of drug candidates [33,34]. Thanks in part to the deluge of molecular data available, recent years have witnessed the adoption of machine learning methods trained on molecular fingerprints or learned representations of chemical structures, tasked with predicting molecular activity in a variety of biological contexts [35–38]. Several of these methods depart from traditional target-oriented approaches to drug discovery, in favour of target-agnostic strategies [39] that employ various phenotypic measurements for model training. Such target-agnostic approach offers new avenues to expand the range of chemical starting points in early phases of the drug discovery pipeline [40], and is particularly well-suited for senolytics discovery given the relatively poor grasp of the molecular pathways that control the senescent phenotype.

Here, we report the development and validation of a machine learning pipeline for the discovery of senolytics. We assembled a dataset mined from multiple sources [11–19,21–26], including academic publications and a commercial patent, and employed it to train machine models predictive of senolytic action. We computational screened a library of more than 5,000 compounds and identified a reduced set of 21 hits for experimental validation. Our experimental screen in two model cell lines of oncogene- and therapy-induced senescence revealed senolytic activity of three compounds: ginkgetin, oleandrin, and periplocin, with potencies and dose-responses comparable to known senolytics. This approach demonstrates that community and open data can be maximally leveraged to detect new active therapeutic compounds, and thus sets the methodological groundwork for a new open science approach to drug discovery and repurposing.

## Results

### Data assembly and quality control

We first assembled a dataset of senolytics (positives) and non-senolytics (negatives) to be used for model training (Fig. 1A). To this end, we mined a panel of 58 senolytics reported in the literature, including compounds from various chemical families such as flavonoids, cardiac glycosides, and antibiotics shown to display senolytic action. These positives have been shown to preferentially target the senescent phenotype in a variety of cell types (Fig. 1B). Some of the compounds, e.g. ouabain [13], have been shown to have wide-spectrum senolytic action, whereas others such as BIX-01294, limit their effect to specific conditions [13]. The literature sources used to assemble the list of 58 senolytics used different cell types, screening assays, and methods for induction of the senescent phenotype. We combined this list with another panel of 21 senolytics reported in a commercial patent [14]. The positives were then merged with a large background of compounds assumed to lack senolytic action. This assumption was needed due to the lack of data on negative screening results from published sources. Since machine learning models can bias their predictions towards the training data they have been exposed to, we chose the negatives from two diverse chemical libraries, LOPAC-1280 and Prestwick FDA-approved-1280, which contain a wide range of FDA-approved or clinical-stage compounds.

**Figure 1.**
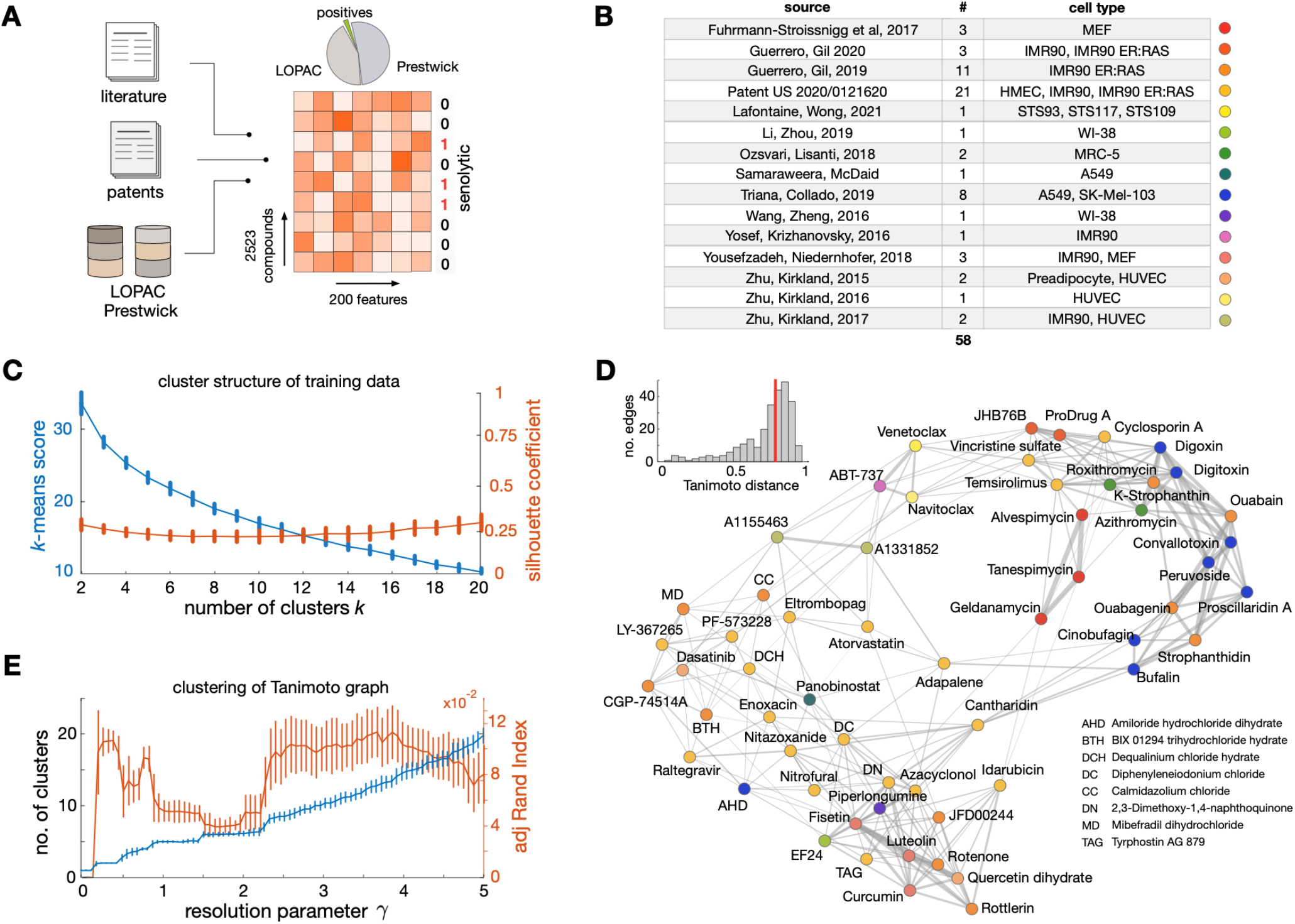
Compounds employed to train machine learning models of senolytic action. **(A)** We assembled a training dataset from multiple sources. We mined 58 known senolytics (positives) from academic papers and commercial patents, and integrated them with a diverse set of compounds drawn from the LOPAC-1280 and Prestwick FDA-approved-1280 chemical libraries assumed to be non-senolytic (negatives). Chemical structures were featurised with 200 physicochemical descriptors computed with the RDKit software [41] and labelled according to their senolytic action. These labelled data were employed to train binary classifiers predictive of senolytic activity. **(B)** Sources of the 58 known senolytics employed for training, including the number of compounds per source and the cell lines where senolytic activity was identified. **(C)** Cluster structure of the senolytics employed for training using the RDKit descriptors as features. Plot shows the *k*-means clustering score and silhouette coefficient [42] averaged across all compounds for an increasing value number of clusters (*k*). Error bars denote one standard deviation over 100 repeats with different initial seeds. The lack of a clear “elbow” in the *k*-means score and low silhouette coefficients suggest poor clustering among the senolytics employed for training. **(D)** Tanimoto distance graph of senolytics employed for training; nodes are compounds and edges represent compounds that are sufficiently close in the RDkit feature space. Node colour indicates the data source as in panel B. To emphasise the overall dissimilarity between compounds, we set the edge thickness as the Tanimoto similarity between compounds (1-distance). Inset shows the distribution of Tanimoto distances across the 269 edges of the graph, with a median distance of 0.77. **(E)** Clustering of the Tanimoto distance graph using the Louvain algorithm for community detection [44]. Plot shows the average number of clusters with respect to the resolution parameter (γ) across 100 runs of the Louvain method (error bars denote one standard deviation); increasing values of y produce a larger number of clusters. We observe pronounced plateaus at 5 and 6 clusters, suggesting some degree of stable clustering in the data. We computed the adjusted Rand index [45] (ARI) averaged across all compounds to quantify the similarity between the Louvain clusters and a labelling obtained from the literature source of each compound (15 labels as in panel E). Low ARI values indicate that Louvain clusters are substantially different from the literature labels.

The full dataset for model training contains 2,523 compounds, including 58 positives (2.3%). We deliberately chose to overrepresent the negatives in the training data so as to reflect the low likelihood of a chemical structure being a senolytic (Fig. 1A). To convert the chemical structures into a numerical format that can be used for model training, we binarized each compound in the training library as 0 (negative) or 1 (positive), and computed 200 physicochemical descriptors with the RDKit package [41]. These descriptors include basic molecular properties such as maximum partial charge, molecular weight, and number of valence electrons, as well as structural properties such as the molecular connectivity Chi indexes, E-State topological parameters, and Kappa shape indexes.

The positive compounds were mined from highly heterogeneous sources that utilised different cell lines and screening assays (Fig. 1B). This bears the risk of introducing bias in our models and limit their predictive power if specific chemical structures or classes are overrepresented in the training data. This is further exacerbated by the heavy imbalance between the number of positives and negatives included in the training data. We thus sought to carefully quantify the diversity of the 58 positives using the RDKit descriptors as feature vectors associated with each compound. To assess diversity of the training data, we examined the cluster structure of the positive compounds using three different methodologies (Fig. 1C-E). We first clustered the positives with the *k*-means algorithm [42] and the cosine distance between feature vectors; this analysis revealed an almost linear decrease in the *k*-means score with respect to the number of clusters (Fig. 1C). The lack of a clear “elbow” in the *k*-means score suggests poor data clustering and hence provides a qualitative indication of diversity in the training data. To determine the quality and consistency of these clusters, we computed the silhouette coefficients for all compounds and number of clusters (*k*) [42]; we found consistently low values for the silhouette coefficient averaged across all compounds, which further suggests little similarity among the senolytic compounds chosen for training.

As a separate check for the diversity of the training data, we built the Tanimoto distance graph for all senolytics employed for training and labelled each compound according to the source from which they were obtained (Fig. 1D). Nodes in the Tanimoto distance graph represent compounds, and two compounds are connected by an edge if they are sufficiently close in the RDkit chemical descriptor space. The structure of the resulting distance graph corroborates the finding that most senolytics are far apart in the descriptor space (median Tanimoto distance = 0.77; Fig. 1D inset), and thus tend to be highly dissimilar to each other. As a final check, we clustered the Tanimoto distance graph using community detection, a type of clustering technique from network science [43] that does not require *a priori* specification of the number of clusters. We employed the popular Louvain algorithm [44], because of its computational efficiency and the inclusion of a resolution parameter (γ) for tuning the granularity of the resulting clusters; larger values of γ lead to more clusters and hence a more granular partition of the graph. For a wide sweep of the resolution parameter, we found an almost linear increase in the number of clusters, and two plateaus at *k*=5 and *k*=6 clusters (Fig. 1E); such plateaus suggest that the data may naturally cluster into five or six groups. We further investigated these clusters and reasoned that compounds may aggregate according to the source where they were mined from. To test this hypothesis, we quantified the similarity between the Louvain clusters and literature labels (Fig. 1B) using the adjusted Rand index (ARI), a useful metric for comparing different clusters that corrects for random group assignments [45]. We found extremely low ARI scores (Fig. 1E) across all cluster resolutions described by the γ parameter; moreover, the ARI scores showed pronounced troughs at the plateaus detected with the Louvain method (mean ARI<0.05 for 100 runs of the clustering method), which we regarded as sufficient evidence that compounds do not cluster according to the source from which they were obtained.

### Predicting senolytic compounds by computational screen using machine learning

We next sought to train machine learning models on the assembled dataset, with the aim of using them to computationally screen chemical libraries and identify hits for experimental validation (Fig. 2A). A key challenge for model training is the heavy imbalance between the number of senolytic and non-senolytic compounds. We thus scored the models with a suite of performance metrics: accuracy (fraction of all compounds correctly identified), precision (fraction of true positive identifications out of all positive identifications), and recall (fraction of correct identifications of true positives). The precision and recall metrics are of paramount importance in our problem, because the class imbalance tends to produce overoptimistic accuracy as a result of correct classification of the majority class, even when the minority class is poorly classified [46].

**Figure 2.**
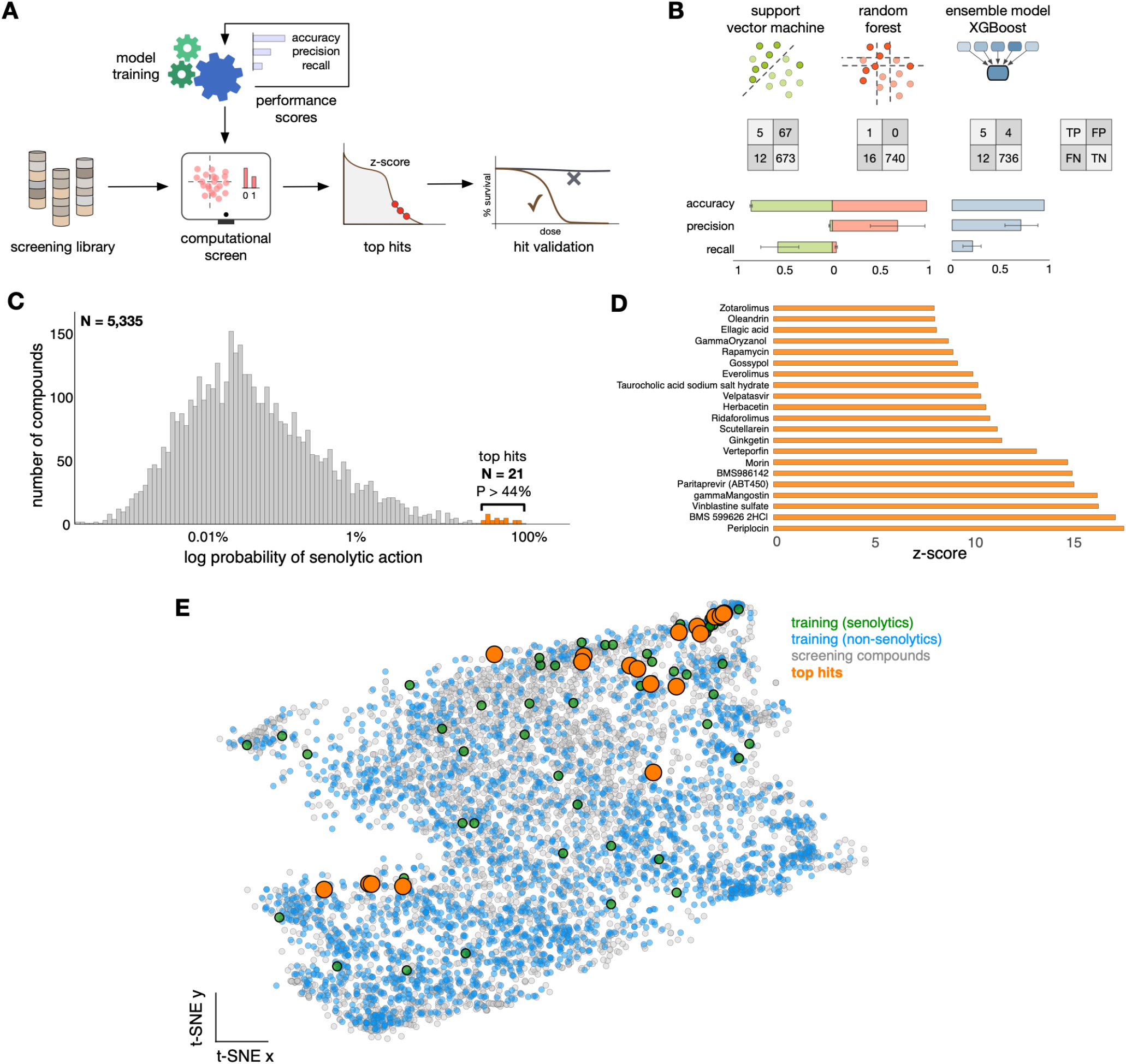
Training of machine learning models and computational screening. **(A)** Pipeline for model training, compound screening, and hit validation. Several classification scores were used as performance metrics to determine the most suitable model for the computational screen. **(B)** Results from three machine learning models trained on 2,537 compounds (Figure 1A); bar plots show average performance metrics computed in 5-fold cross validation, with error bars denoting one standard deviation across folds. The confusion matrices were computed from models trained on 70% of compounds, and tested on 20 positives and 742 negatives that were held-out from training. All models displayed poor performance metrics (Supplementary Table 1), and we chose the XGBoost algorithm for screening because of its lower number of false positives; the high accuracy of all models is a well-known artefact in classification problems with heavy class imbalance. **(C)** Results from computational screen of the L2100 TargetMol Anticancer and L3800 Selleck FDA-approved & Passed Phase chemical libraries, totaling 5,335 compounds. The XGBoost model is highly selective and scores the vast majority of compounds with an extremely low probability of having senolytic action; a small fraction of N=21 compounds were scored with P>44%, which we selected for experimental validation. **(D)** Compounds selected for screening, ranked according to their z-score normalised prediction scores from the XGBoost model; the selected compounds are far outliers in the distribution of panel C. **(E)** Two dimensional t-SNE visualisation of all compounds employed in this work; t-SNE plots were generated with perplexity = 50, learning rate 200, and maximal number of iterations 1,200 [48]. Predictions scores above 44% from the XGBoost model are marked with orange circles.

We focused primarily on two common models for binary classification: support vector machines (SVM) and random forests (RF). These models operate by partitioning the feature space so as to ensure that positive and negative samples are optimally assigned to a partition. In their basic form, SVM slices the feature space with a hyperplane, while RF are ensembles of decision trees that segment the feature space with orthogonal cuts across each feature [42]. Models were trained using gradient-based optimization on a reduced set of 165 normalised RDKit features that were identified with feature importance analysis (Supplementary Fig. 1A), and using k-fold cross-validation to avoid overfitting and robust model comparison.

We found that both SVM and RF models displayed poor performance (Fig. 2B) but showed marked differences in the type of misclassification errors they produce. The confusion matrices (Fig. 2B) suggest that the RF model tends to return few false positives (high precision) and a high number of false negatives (low recall), whereas the SVM returns opposite results. We observed similar trade-offs between false negatives and false positives in various other models of varied complexity (Supplementary Table 1), including simple logistic regressors, Bayesian classifiers, and deep learning models [37]. In line with expectation, the majority of models displayed high accuracy (>90%) but this is an artefact resulting from the correct classification of many non-senolytic compounds that are overrepresented in the training data.

For the purposes of early-stage drug discovery, false positives are more deleterious than false negatives because they artificially inflate the number of predicted hits and thus increase the costs of downstream experimental validation. We thus took the performance of the RF model as a baseline, and aimed at improving its predictive power by using an ensemble model (XGBoost) that is known to improve performance by aggregating predictions from a collection of decision trees [47]. The XGBoost model improved both precision and recall scores, and overall returned the best performance among all considered models (Fig. 2B). We observed precision scores of up to 75% in some of the cross-validation runs, which amounts to a false discovery rate of 25% that we regarded as acceptable given the heterogeneity of the data employed for training.

We next employed the pre-trained XGBoost model to computationally screen a library of chemical structures selected on the basis of chemical diversity. We screened a list of compounds consisting of the contents from the L2100 TargetMol Anticancer and L3800 Selleck FDA-approved & Passed Phase libraries into a single dataset with 5,335 compounds featurised with the physicochemical descriptors from RDKit; none of the compounds in the screening library were employed for training. The computational screen revealed the XGBoost model to be exceptionally selective (Fig. 2C), with the vast majority of compounds having extremely low prediction scores, and thus deemed to have a low probability of being senolytic. Importantly, the score distribution (Fig. 2C) displayed a small group of 21 compounds (0.4% of the full library) with a comparatively higher probability of being senolytic (Fig. 2C, orange); this small set of compounds were scored above 44% by our XGBoost model and were selected for further experimental validation. We note that the selected compounds are extreme outliers, with prediction scores at least 8 standard deviations away from the bulk of compounds screened (Fig. 2D). We employed dimensionality reduction to visualise the training and screening compounds in the RDKit feature space [48], which revealed a strong overlap between the two sets and thus a strong evidence that the computational screen was performed in the high-confidence domain of the machine learning model (Fig. 2E).

### Identification of senolytics by experimental screening of top predicted compounds

To validate our machine learning approach, we experimentally screened the top predicted molecules (Fig. 2D) for senolytic activity in two model cell lines for oncogene-induced and therapy-induced senescence. We first assessed oncogene-induced senescence (OIS) in human diploid fibroblasts IMR90 transduced with the fusion protein ER:RAS (IMR90 ER:RAS), which induces oncogenic Ras^G12V^-mediated stress by addition of 4-hydroxytamoxifen (4-OHT) to the culture media [49]. Treatment of IMR90 ER:RAS cells with 4-OHT showed a decrease in proliferation, increased senescence associated β-galactosidase activity, induction of cell cycle inhibitor expression, and activation of the SASP when compared with control and 4-OHT untreated cells, indicating that the cells underwent oncogene-induced senescence (Supplementary Fig. 2).

To test the senolytic activity, we compared the effect of each compound on the total cell number (automated high content image-based analysis of total number of nuclei per well) in non-senescent and senescent IMR90 ER:RAS cultures treated with the top hits from the computational screen (Fig. 2D, Fig. 3A-D, Supplementary Fig. 3A-C). The drop in cell number compared to untreated controls is indicated by the nuclei count and reflects cell death [13,15]. As a positive control, we used ouabain, a cardiac glycoside with well characterised senolytic activity [13]. An optimal senolytic effect was reached with addition of 46.4 nM ouabain (IC50 control = 231 nM; IC50 senescence = 28 nM) to 4-OHT-induced IMR90 ER:RAS cells. This concentration killed most of the cultured senescent cells, but resulted in no reduction in numbers of non-senescent control cells (Fig. 3B).

**Figure 3.**
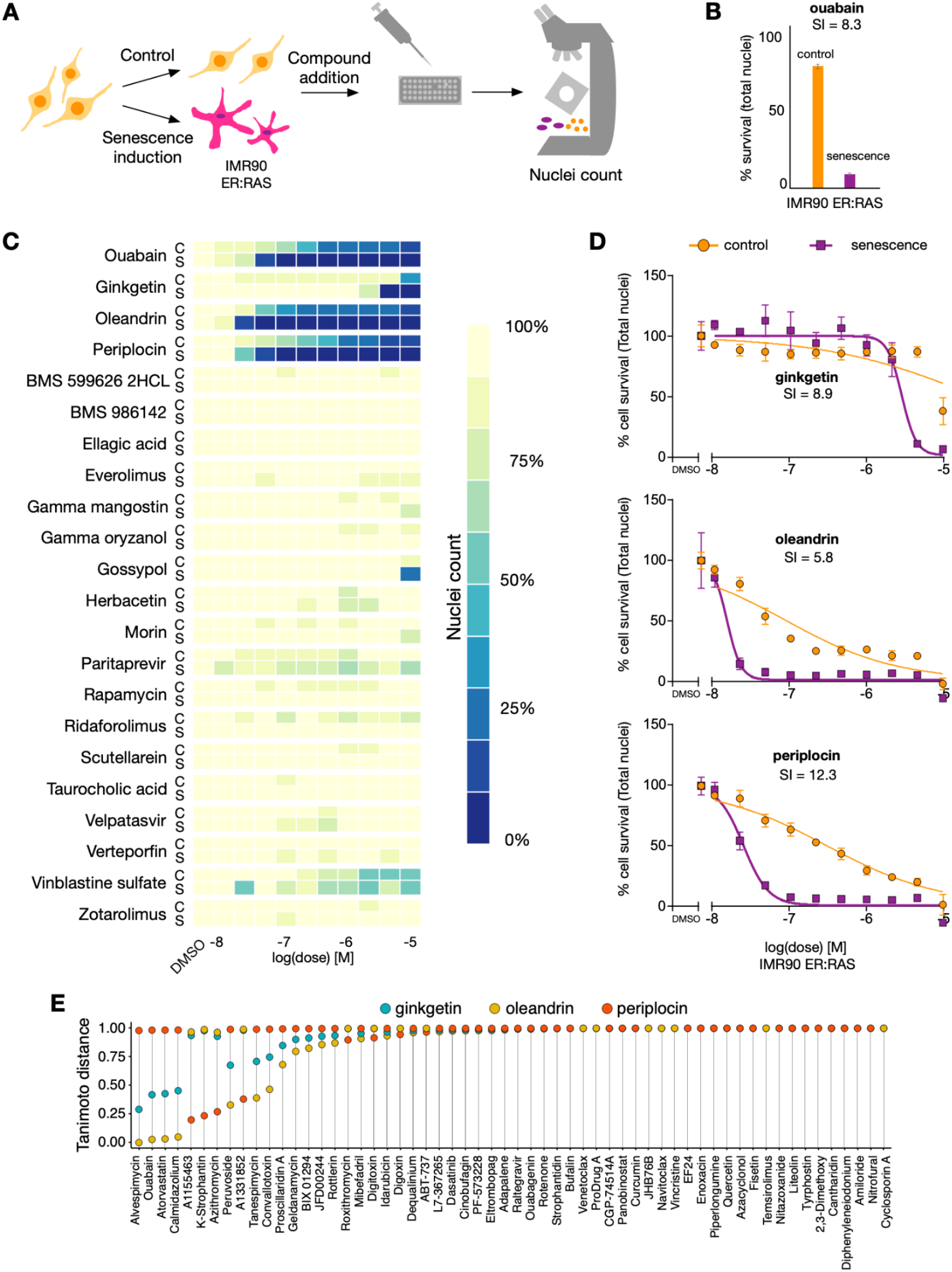
Experimental characterisation of compounds selected for screening in oncogene-induced senescent cells. **(A)** Experimental setup of oncogene-induced senescence model with IMR90 ER RAS cells. Senescence was induced by addition of 4-OHT at 100 nM during thet duration of the experiment (eight days). Control and senescent cells were plated in a 96 well plate day five of 4-OHT induction. Top predicted compounds were added after multiwell seeding, and 72 hours afterwards, the cells were fixed, and the nuclei stained and counted. **(B)** Bar plot of OIS positive experimental control, ouabain, at 46.4 nM. Data is normalised to DMSO. Mean ± s.d. are shown from n=3 experiments. **(C)** Results from experimental validation of controls and the top 21 compounds from Fig. 2D predicted to have senolytic action with P>44%. Three compounds out of the 21 displayed senolytic activity: ginkgetin, oleandrin, and periplocin. **(D)** Dose-response curves of the three newly found senolytic compounds. The senolytic index (SI) is defined as the ratio between the IC50 of control cells and the IC50 of senescent cells. Data is normalised to DMSO. Mean ± s.d. are shown from n=3 experiments. (**E**) Tanimoto distance from each of the three validated senolytics to each senolytic employed for training; distances were calculated using the RDKit descriptors that were employed in the training of machine learning models in Fig. 2B and Supplementary Table 1.

We then tested the 21 candidate compounds and found three with clear senolytic action: periplocin and oleandrin, two cardiac glycosides which have not been previously identified as senolytics, and ginkgetin, a natural non-toxic biflavone; this amounts to a hit confirmation rate of 14.28% (Fig. 3C-D, Supplementary Fig. 3C). Treatment of senescent IMR90 ER:RAS cells with the three compounds showed reduced nuclei counts when compared with proliferating, non-senescent IMR90 controls with an effect comparable to the positive control (Fig. 3C-D, Supplementary Fig. 4A). For confirmation, we performed cell staining with Hoechst to label the nuclei, showing the clearest effect with doses of 21.5 nM oleandrin (IC50 control = 85 nM; IC50 senescence = 14 nM), 46.4 nM periplocin (IC50 control = 300 nM; IC50 senescence = 24 nM) and 4.6 µM ginkgetin (IC50 control = 26 µM; IC50 senescence = 2.6 µM). These concentrations have a marginal effect on normal cells’ nuclei counts, but a marked decrease in nuclei counts in senescent cells (Fig. 3C-D, Supplementary Fig. 4A). The dose-response curves showed a lower IC50 for the three compounds in nuclei counts from senescent cells as compared to nuclei from normal cells (Fig. 3D). In particular, periplocin (the top hit predicted by our model, Fig. 2D) showed a senolytic effect fully comparable to the positive control ouabain (Fig. 3B and Fig. 3D).

We performed a second validation of the effectiveness of our machine learning models using a different stressor. We focussed on therapy-induced senescence (TIS, Supplementary Fig. 5A), where human epithelial cancerous cells (A549) were induced to become senescent by addition of etoposide [49]. Cells were treated with 100 µM etoposide for 48 hours, followed by another three days of exposure to media in standard conditions. As a positive control, we employed navitoclax (IC50 control = 10.2 µM; IC50 senescence = 440 nM), a Bcl-2 family inhibitor with well characterised senolytic activity [26]. Addition of 1 µM navitoclax to A549 cells killed most of the cultured senescent cells, but resulted in no reduction in numbers of non-senescent control cells, confirming its optimal senolytic activity (Supplementary Fig. 5B).

Cells were then treated with the top 21 hits in Fig. 2D from our computational screen (Supplementary Fig. 5A-D, Supplementary Fig. 6A); we found that the same three compounds validated in oncogene-induced senescent cells (periplocin, oleandrin, ginkgetin) also displayed strong senolytic action in senescent A549 cells. The three compounds showed enhanced toxicity when compared with proliferating, non-senescent A549 controls with an effect comparable to the positive control. Hoechst labelling showed that a senolytic effect was reached with concentrations of 10 nM oleandrin (IC50 control = 19.5 nM; IC50 senescence = 5.4 nM), 46.4 nM periplocin (IC50 control = 267 nM; IC50 senescence = 72.2 nM) and 4.64 µM ginkgetin (IC50 control = 10.4 µM; IC50 senescence = 5.7 µM). These doses have a marginal effect on normal cells, but a decrease in survival rate in senescent cells (Supplementary Fig. 4B, Supplementary Fig. 5C-D). The dose-response curves showed a lowered IC50 for the three compounds in senescent cells as compared to normal cells (Supplementary Fig. 5D).

To assess the similarity between the three compounds correctly identified by the XGBoost model (Fig. 3D) and the senolytics employed for training (Fig. 2B), we computed the Tanimoto distance in the descriptor space between ginkgetin, oleandrin, and periplocin and each of the 58 senolytics in the training data (Fig. 3E). More than half of the training compounds were found to be maximally distant from our newly discovered senolytics, which provides strong validation that our machine learning approach can effectively identify diverse compounds for specific biological effects such as senolysis.

## Discussion

Current approaches to drug discovery suffer from notoriously high attrition rates in late-stage preclinical and clinical development. Due to their ability to parse and detect patterns in large volumes of data, AI has found applications across every stage of the drug discovery pipeline [50]. In this paper, we described a successful machine learning approach designed to identify novel drug candidates in early phases of the discovery process. We focused on targeted elimination of senescent cells, a phenotype that has attracted substantial interest for adjuvant cancer therapy [2], but for which few molecular targets have been identified. Our strategy revealed three compounds (ginkgetin, oleandrin, periplocin) that selectively eliminate cells displaying oncogene- and therapy-induced senescence. We showed that these compounds have a potency comparable to senolytics previously described in the literature and, crucially, our method led to large gains in efficiency by reducing the number of compounds for experimental screening by more than 200-fold.

Our approach offers several innovations that depart from current practice in AI for drug discovery. First, it relies solely on published data for model training, and thus avoids the extra costs for in-house experimental characterisation of training compounds. Second, our machine learning models were trained on just 58 chemical structures with proven senolytic action, which is much smaller data than typically considered in the field; the size of our training set is a direct consequence of senolysis being a rare molecular property and the limited number of senolytics reported in the literature so far. The success of our approach demonstrates that machine learning can take maximum advantage of literature data, even when such data is heterogeneous and of much smaller scale than typically expected [51]. Third, our models were trained in a target-agnostic manner using phenotypic signatures of drug action. Target specificity is of key importance for drug efficacy and safety in later stages of the discovery pipeline, but there are numerous conditions of high economic and societal burden with few or no known targets [40]; for such conditions, there is an opportunity for phenotypic drug discovery to increase the number of chemical starting points that can be carried through the discovery pipeline [39].

A key challenge in computational drug screening is the construction of numerical representations of chemical structures that are predictive of drug efficacy [52]. With the advent of deep learning as the leading paradigm in the field [53], many recent works have developed such representations with e.g. transformer models for prediction of chemical reactions [54], graph neural networks to describe molecular structures [37,38], and generative models for *de novo* compound design [55]. In our approach, we found that classic physicochemical descriptors [41] were sufficient to train useful models and, moreover, we observed limited benefits in the use of deep learning for compound featurisation (Supplementary Table 1), possibly as a result of the small size of our training data. We instead found that careful data assembly, curation, and quality control were the key for success. Since negative assays are rarely reported in the literature, we built the training data by pairing the known senolytics with a background of compounds assumed to lack senolytic action, but with an appropriate chemical diversity and a size deliberately chosen to reflect the paucity of senolytic compounds. These design choices caused a heavy imbalance between the number of senolytic and non-senolytic compounds, which introduced additional challenges for model training. Several checks were needed to ensure that the training data was diverse enough and avoided bias toward specific chemical classes. Moreover, our models generally displayed poor performance as quantified by common classification metrics, producing large numbers of false positives and false negatives in our cross-validation analyses. We mitigated the impact of class imbalance by prioritising models with a lower number of false positives, so as to reduce the downstream costs for experimental validation. Our results thus show that seemingly poor models can still be effective with adequate checks and balances on the structure of the training data, plus a careful interpretation of performance metrics and misclassification errors.

Our approach led to a significant reduction in experimental screening costs, largely because all models were trained solely on published data and, unlike other recent successes in the field [37], there was no need to screen compounds purposely for model training. The approach thus offers exciting prospects for new open science approaches to drug discovery. Most recently, the COVID-19 pandemic spurred a multitude of such initiatives across the globe with the goal of finding new antivirals from the troves of published data [56]. Our work provides a concrete example of a simple yet effective machine learning pipeline that can be readily built from published screening data. We hope this approach will catalyse more open science approaches to discover treatments for conditions of unmet need, particularly those for which there is a limited grasp of the biological pathways involved in disease onset and progression.

## Methods and Materials

### A Data assembly, featurisation and quality control

#### Training data

We assembled a list of 58 previously identified senolytics mined from 15 sources [11–19,21–26] (Fig. 1B). The library of negative compounds contains 2,465 compounds from the LOPAC-1280 (Library of Pharmacologically Active Compounds; Merck, Darmstadt, Germany) and Prestwick FDA-approved-1280 (Prestwick Chemical, Illkirch, France). We reasoned that these libraries are sufficiently diverse for training machine learning models. Although all the compounds from these two library sources were deemed as negative, there is a possibility that some of these molecules have been incorrectly labelled. This is because not all senolytics found have been expressly named in all high-throughput publications (some sources only name a small set of their discoveries) or because these molecules have only been tested in one or two cell lines under one type of senolytic induction process.

#### Featurisation

The training dataset contains a one-dimensional vectorial representation of the two-dimensional structure of every molecule in the form of a SMILES string. The majority of SMILES were taken from the library of origin of every compound (LOPAC or Prestwick for training, Selleck or TargetMol for screening) with the exception of five positives (ProDrug A, JHB76B, CGP-74514A, A1331852, A1155463) which SMILES were calculated using ChemDraw [12,13,19]. For chiral molecules, we favoured isomeric SMILES representations instead of the canonical case. We employed the RDKit package [41] to compute 200 physicochemical descriptions for each compound, which quantify different aspects of the molecular structure, its fragments, and global characteristics.

#### Clustering analysis and Tanimoto graph

To quantify the diversity of the senolytics employed for training (Fig. 1C-E), we performed *k*-means clustering of the 58 positives using the RDKit descriptors as feature vectors and cosine distance function between z-score normalised feature vectors. The degree of clustering was quantified by the *k*-means score (Fig. 1C), i.e. the optimal value of the loss function employed to determine the clustering. The quality of the *k*-means clusters were determined with the silhouette coefficient *S* averaged across the 58 senolytics. The silhouette coefficient varies between −1 and 1, with *S*=1 indicating that compounds are in well separated clusters, *S*=0 indicating overlapping clusters, and *S*=-1 indicating incorrect assignment of clusters. To build the Tanimoto distance graph (Fig. 1D) we first constructed a fully connected graph weighted by the pairwise Tanimoto distances feature vectors. This graph was then sparsified with the *k*-nearest neighbours graph (*k*=7) intersected with the minimum spanning tree of the original graph. The edge widths were set as the Tanimoto similarity between compounds (1-distance). Clustering of the Tanimoto distance graph (Fig. 1E) was done with a Matlab implementation of the Louvain algorithm for community detection [44]. To compare the Louvain clusters with the compounds labelled according to their source (Fig. 1B), we employed the adjusted Rand Index which is a measure of the similarity between two clusterings, adjusted for the chance grouping of compounds [45]; low values of the ARI indicate little similarity between clusterings.

### B Model training and computational screen

All models were trained with the scikit-learn 0.24.1 Python library, plus the XGBoost v 0.90 [47]. Models were trained on a reduced set of 165 z-score normalised features identified as relevant for classification using scikit-learn feature importance with a forest of trees function (Supplementary Fig. 1). All models were scored with three metrics of classification performance:

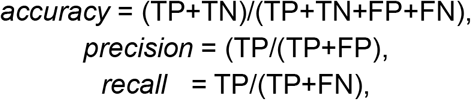

where TP, TN, FP and FN are the number of true positives, true negatives, false positives and false negatives, respectively. Models were trained and evaluated with a random 70:30 split of the data (70% for training, and 30% for testing). We stratified samples in the random split to ensure that both the positive and negative classes were proportionally represented in the training and testing sets. In all cases, a 5-fold cross-validation was performed. For the final XGBoost model employed for the computational screen, we optimised the hyperparameter tree-depth and obtained consistent performance over 5-fold cross-validation runs, with values varied from 1 to 11; we settled tree-depth = 10 to allow for a more profound use of our high-dimensional feature space. For the computational screen, we used the L2100 TargetMol Anticancer (TargetMol Chemicals, Wellesley Hills, MA) and L3800 Selleck FDA-approved & Passed Phase (Selleck Chemicals, Houston, TX) libraries due to the versatility of compounds represented in them. We used the pre-trained XGBoost model to compute prediction scores on the augmented library, i.e. the probability of a compound being classified as senolytic (Fig. 2C).

### C Validation assays

We performed experimental validation of the compounds with z-score>8 (Fig. 2D). These correspond to 21 compounds out of a total of 5,335 compounds. We employed two cellular models of senescence one of OIS and a second one of TIS. For the OIS case, we utilised IMR90 ER:RAS cells with 4-OHT at 100 nM. The 4-OHT treatment had a duration of six days. For the TIS model, we used A549 cells with etoposide at 100 µM. The exposure of the cells to etoposide lasted 48 hours, after which period the cells were cultivated for a further three days with normal media.

#### Cell culture

IMR90 female human foetal lung fibroblasts and A549 human lung adenocarcinoma cells were obtained from the American Type Culture Collection (ATCC, Manassas, VA). The cell lines were confirmed to be mycoplasma negative (Lonza MycoAlert, cat #LT07-118). IMR90 ER:RAS is a derivative of IMR90 cells expressing a switchable version of oncogenic H-Ras [57]. IMR90 ER:RAS and A549 cells were maintained in Dulbecco’s Modified Eagle Medium (DMEM, ThermoFisher) supplemented with foetal bovine serum (10%), L-glutamine (2 mM, ThermoFisher), and antibiotic-antimycotic solution (1%, ThermoFisher) and incubated under standard tissue culture conditions (37 °C and 5% CO2).

#### Quantification of senolytic action

Cells were seeded (50 µL per well) into 384-well plates (IMR90 ER:RAS cells in Nunc Optical Bottom Polybase Microplates [#142761, Thermo Scientific, Rochester, NY] and A549 cells in CELLSTAR Cell Culture Microplates [#781091, Greiner Bio-One, Kremsmunster, Austria]). Cells were incubated under standard tissue culture conditions for 24 h before the addition of compounds. Passage 14 IMR90 ER:RAS cells were seeded at 1,300 cells per well in the control condition, and at 1,600 cells per well in the senescent case. Passage 34 A549 cells were seeded at 7,000 cells per well in the control condition, and at 10,000 cells per well in the senescent case.

Dose response plates were prepared with a DMSO control and a 10-point half-log concentration range, and added to the compounds using a D300e digital dispenser (Tecan Trading AG, Switzerland) at a final concentration of between 10 uM and 10 nM. Every screened condition was carried out in triplicate. After 72h of incubation with exposure to the compounds, cells were fixed by the addition of an equal volume of formaldehyde (8%, 50 uL; #BP531-500, Fisher Bioreagents, Fisher Scientific, Loughborough, Leicestershire) to the existing media, incubated at room temperature (30 min), and washed three times in phosphate-buffered saline (PBS). Cells were then permeabilized in Triton-X100 (0.1%, 50 µL) and incubated at room temperature (30 min) followed by three more washes with PBS.

Cells were stained with Hoechst 33342 for nuclei count (excitation/emission wavelength at 387l447 nM, DAPI channel, original concentration at 10 mg/ml, final concentration at 2 µg/ml; H1399, Molecular Probes, Eugene, OR). The staining solution was prepared in bovine serum albumin solution (10 %). The staining solution was added to each well (30 µL) and incubated in the dark at room temperature (30 min), followed by three washes with PBS and no final aspiration. Plates were foil sealed.

#### Image acquisition

Plates were imaged on an ImageXpress micro XLS (Molecular Devices, Eugene, OR) equipped with a robotic plate loader (Scara4, PAA, UK). Four fields of view were captured per well (20x objective for A549 cells, 10x objective for IMR90 ER:RAS cells) and one filter was used (DAPI). A typical wild-type field of view contained 1000 cells in the IMR90 ER:RAS case, and 1400 in the A549 case.

#### Image and data analysis

The stained cell nuclei were counted on MetaXpress v6.6.2.46 software. The results per compound, phenotype condition, and dose were added and the results morphed into data frame format with functions from R’s dplyr, tidyr, and reshape2 libraries.

Concentration response data (control plus 10 half-log range points) was fitted (ordinary least squares) to a log(inhibitor) vs normalised response (control value per condition [senescent, non-senescent] was constrained at 100%) with variable slope equation using Prism v6 software (GraphPad, San Diego, CA). With this fit, IC50 values were calculated for senolytic compounds.

#### Compounds

The following compounds were used in the present study: etoposide (Sigma-Aldrich, E1383), 4-hydroxytamoxifen (Sigma-Aldrich, H7904), ouabain (Apexbio, B2270), navitoclax (Apexbio, A3007), ginkgetin (Cayman Chemical, 25103-1mg), oleandrin (Cayman Chemical, 29871-1mg), periplocin (Cayman Chemical, 25216-1mg), BMS 599626 dihydrochloride (Apexbio B5792), BMS 986142 (BioVision, B2420-1), ellagic acid (Apexbio, A2306), everolimus (Cayman Chemical, 22559-1mg), herbacetin (Cayman Chemical 19285-1mg), morin (MedChemExpress LLC, HY-N0621-10mg), paritaprevir (MedChemExpress HY-12594), rapamycin (Cayman Chemical, 13346-5mg), taurocholic acid sodium salt hydrate (Selleck Chemicals, S5130), velpatasvir (BioVision, B1194-5), verteporfin (Apexbio, A8327), zotarolimus (Cayman Chemical, 29246-5mg), gamma mangostin (MedChemExpress LLC, HY-N1957-5mg), gamma oryzanol (MedChemExpress LLC, HY-B2194), gossypol (MedChemExpress LLC, HY-15464A-10mg), ridaforolimus (Apexbio, B1639), scutellarein (MedChemExpress LLC, HY-N4314-1mg), vinblastine sulfate (MP Biomedicals LLC, 0219028725).

## Acknowledgements

The authors disclosed receipt of the following financial support for the research, autorship, and/or publication of this article: VSB is a cross-disciplinary post-doctoral fellow supported by funding from the University of Edinburgh and Medical Research Council (MC_UU_00009/2). JCA acknowledges funding by Cancer Research UK (CRUK) (C47559/A16243 Training & Career Development Board - Career Development Fellowship), the University of Edinburgh Chancellor’s Fellowship R42576 MRC, the Ministry of Science and Innovation of the Government of Spain (Proyecto PID2020-117860GB-I00 financiado por MCIN/ AEI /10.13039/501100011033) and the Spanish National Research Council (CSIC).

