## Supplementary Material for "Discovery of new senolytics using machine learning"

### Supplementary Methods

#### Senescence induction and assays

**Cell culture.** IMR90 human fibroblasts were grown in DMEM (Thermo Fisher Scientific) supplemented with 10% FCS and 1× Anti-Anti (Gibco). Cells were kept in culture at 37 % $CO_2$ and ambient $O_2$ at 37 °C.

**Western Blot analysis.** Cell lysates were prepared using Cell Lysis Buffer (Cell Signalling 9803S). Clear lysates were quantified by Bradford colorimetric assay. Samples were resolved by polyacrylamide gel electrophoresis and transferred on nitrocellulose membrane, which was blocked by a 5% milk-TBS-Tween buffer for 1h at RT. Primary antibodies (anti-p21 Sigma P1484, anti-p16 JC8, anti-IL1B MAB201 R&D) were incubated o/n at 4 °C. After 2 washes with TBS-Tween buffer, secondary antibodies were added for 1h at RT and then 2 washes were done before developing by using enhanced chemical luminescence (Amersham) detection reagent. HRP-B-actin was incubated for normalisation.

**Cell proliferation assay.** 50,000 cells per 10 cm-plate were seeded and kept in culture. After 12 days, cells were fixed with 1% glutaraldehyde (Sigma) for 1h. After several washes with water, dried plates were stained with 0.15% crystal violet solution for 1h and then washed again. Once they were completely dried, plates were scanned for analysis.

**SA-β-galactosidase assay.** After 10 days in culture, cells were fixed in 0.5% glutaraldehyde (Sigma) for 10 min at RT. Then the cells were washed and stained with SA-β-Gal staining solution (20× KC [100 mM K3FE (CN)6 and 100 mM K4Fe(CN)6*3H2O in PBS], 20× X-Gal solution (Thermo Fisher Scientific) diluted to 1× in PBS/1 mM $MgCl_2$ pH 6). Staining was performed overnight at 37 °C in the dark. Once cells were washed, images were taken using an inverted tissue culture widefield microscope (Nikon) for documentation and quantification.

**Immunofluorescence and imaging.** IMR90 ER:STOP and IMR90 ER:RAS cells treated with 4-OHT during 8 days were fixed with 4% paraformaldehyde for 30 min. After several washes, cells were permeabilized with 0,2% Triton-100 for 10 min and then blocked for 30 min with PBS-BSA-Gelatin fish (Sigma). Primary antibodies (IL1A AF-200-NA R&D Systems; IL1B MAB201 R&D Systems; and IL8 MAB208) were prepared in a blocking buffer and incubated for 1h at RT. Alexa Fluor 594 goat anti-mouse was used for signal detection and DAPI solution was added for 10 min. Finally, samples were washed before imaging. Confocal images (512x512 pixels; 0.76um/pixel) were acquired sequentially on a SP5 laser-scan microscope (Leica) with a 20× NA objective and 2× electronic zoom using LAS AF acquisition software. Cells were excited sequentially with 405 nm and 594 nm laser lines and emission was captured between 430-480nm (DAPI) and 605-655nm (Alexa594) respectively. Images are presented after digital adjustment of curve levels to maximise signal with ImageJ software. In all cases, exposure time, sensor gain, and digital manipulation were the same for control and experimental samples. Fluorochromes and colours are as indicated in the figure legends.

### Supplementary Figures and Tables

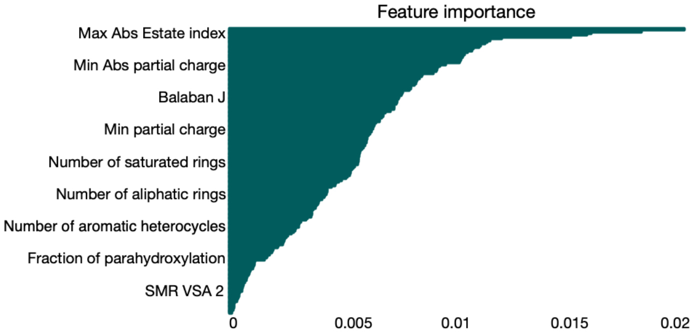


**Supplementary Figure 1. Evaluation of feature importance for training of machine learning models.** Features used for training of machine learning models from RDKit, ranked by feature importance percentage. 165 descriptors out of 200 calculated gave a non-zero importance score, and were therefore utilised to train the machine learning models described in Fig. 2 and Supplementary Table 1. A description of these descriptors may be found on the RDKit website [41].

| Model | Accuracy | Precision | Recall |
|---|---|---|---|
| K-nearest neighbours (K=5) | 0.98 ± 0.00 | 0 | 0 |
| Gaussian Bayes | 0.65 ± 0.08 | 0.06 ± 0.03 | 0.64 ± 0.08 |
| Logistic regression | 0.98 ± 0.00 | 0 | 0 |
| Decision tree | 0.96 ± 0.01 | 0.16 ± 0.05 | 0.22 ± 0.09 |
| Voting classifier (RF, SVM, XGBoost) | 0.98 | 0.63 ± 0.37 | 0.08 ± 0.05 |
| SMOTE with Tomek links | 0.95 ± 0.01 | 0.19 ± 0.06 | 0.35 ± 0.22 |

**Supplementary Table 1. Performance metrics of binary classifiers trained on the assembled dataset of senolytics.** At a preliminary stage, a variety of models were trained for the classification task at hand, including SMOTE, a method specialised for datasets with imbalanced classes [58]. The performance of these models was consistently poor compared to that of the models in the intermediate stage of training (SVM, RF, XGBoost). Additionally, we trained the deep-message passing neural network (D-MPNN) described in Refs. [37,38] with our data, and calculated its performance with the metrics available in the chemprop software and 5-fold cross-validation. A D-MPNN model without additional features displayed accuracy = 0.97 ± 0.01 and precision-recall-auc = 0.25 ± 0.17. Addition of RDKit features to the D-MPNN gave accuracy = 0.97 ± 0.01 and precision-recall-auc = 0.34 ± 0.21.

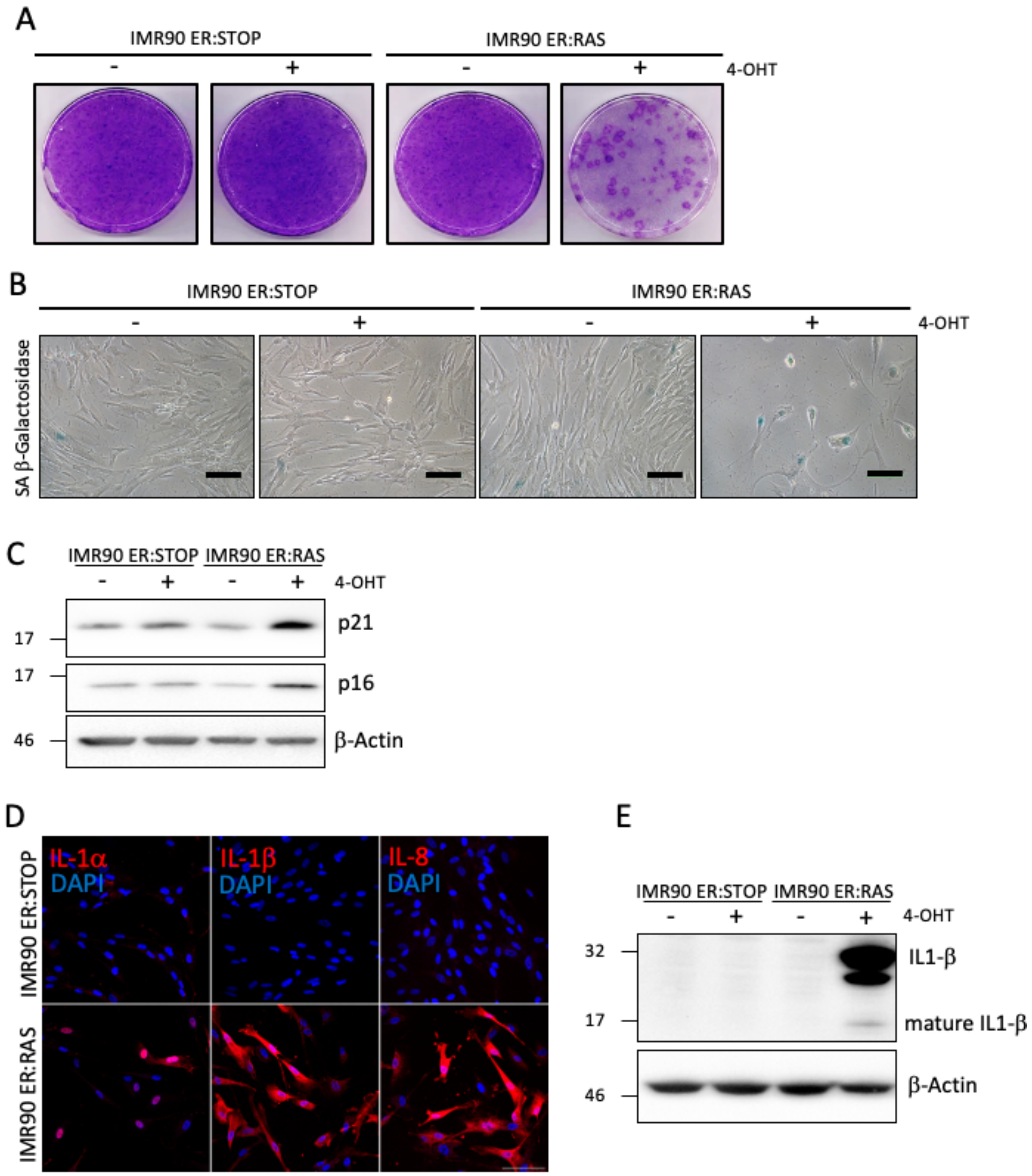


**Supplementary Figure 2. Oncogene-induced senescence model (OIS) characterization in human diploid fibroblasts IMR90. (A)** Same number of IMR90 ER:STOP and ER:RAS cells were seeded with or without 100 nM 4-OHT as indicated and cultured for 12 days and then stained for crystal violet. **(B)** SA-β-galactosidase assay was performed 10 days after 4-OHT treatment. Scale bars, 100 µm. **(C)** Western blot of p21 and p16 expression in IMR90 ER:STOP and IMR90:RAS cells treated with 4-OHT for 4 days. β-actin is shown as loading control. **(D)** Immunofluorescence labelling for Il-1a, IL-1b and IL-8 8 days after 4-OHT treatment. Representative images obtained in a confocal microscope are shown (scale bars, 100 µm). **(E)** Western blot of IL-1b expression in IMR90 ER:STOP and IMR90:RAS cells 8 days after 4-OHT treatment.

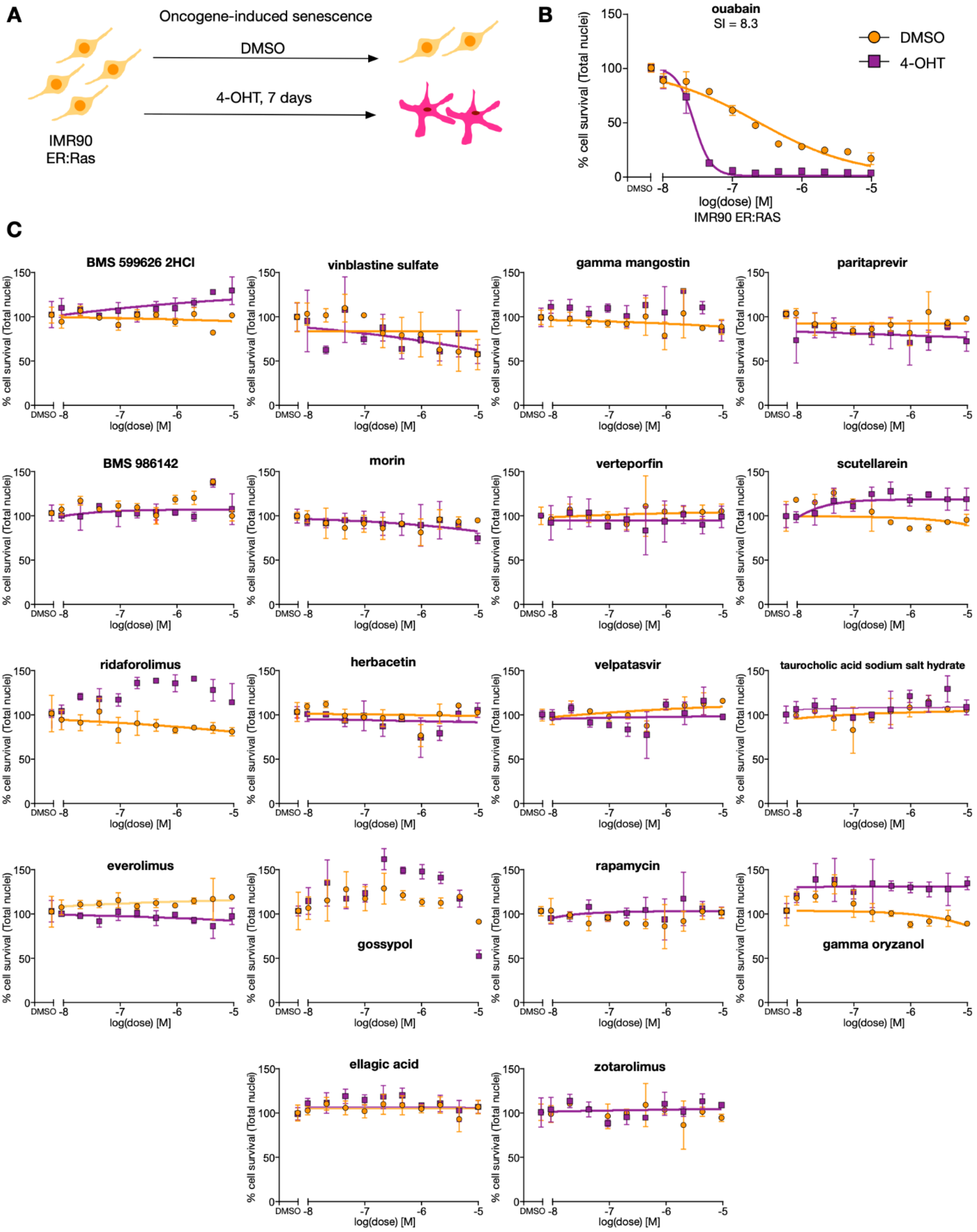
A
Oncogene-induced senescence
DMSO
4-OHT, 7 days
IMR90
ER:Ras
B
ouabain
SI = 8.3
DMSO
4-OHT
% cell survival (Total nuclei)
log(dose) [M]
IMR90 ER:RAS
C
BMS 599626 2HCl
vinblastine sulfate
gamma mangostin
paritaprevir
BMS 986142
morin
verteporfin
scutellarein
ridaforolimus
herbacetin
velpatasvir
taurocholic acid sodium salt hydrate
everolimus
gossypol
rapamycin
gamma oryzanol
ellagic acid
zotarolimus

**Supplementary Figure 3. Experimental characterisation of false positives predicted by the machine learning model in oncogene-induced senescence. (A)** Experimental setup of oncogene-induced senescence model with IMR90 ER:RAS cells. Senescence was induced by addition of 4-OHT at 100 nM for six days. Control and senescent cells were plated on the last day of senescence induction. Top predicted compounds were added a day later, and 72 hours afterwards, the cells were fixed, and the nuclei stained and counted. **(B)** Dose-response curve of OIS positive experimental control, ouabain. Data is normalised to DMSO. Mean ± s.d. are shown from n=3 experiments. **(C)** Dose-response curves of false positives of the XGBoost model (non-senolytic compounds predicted to have senolytic action with P>44%). Data is normalised to DMSO. Mean ± s.d. are shown from n=3 experiments.

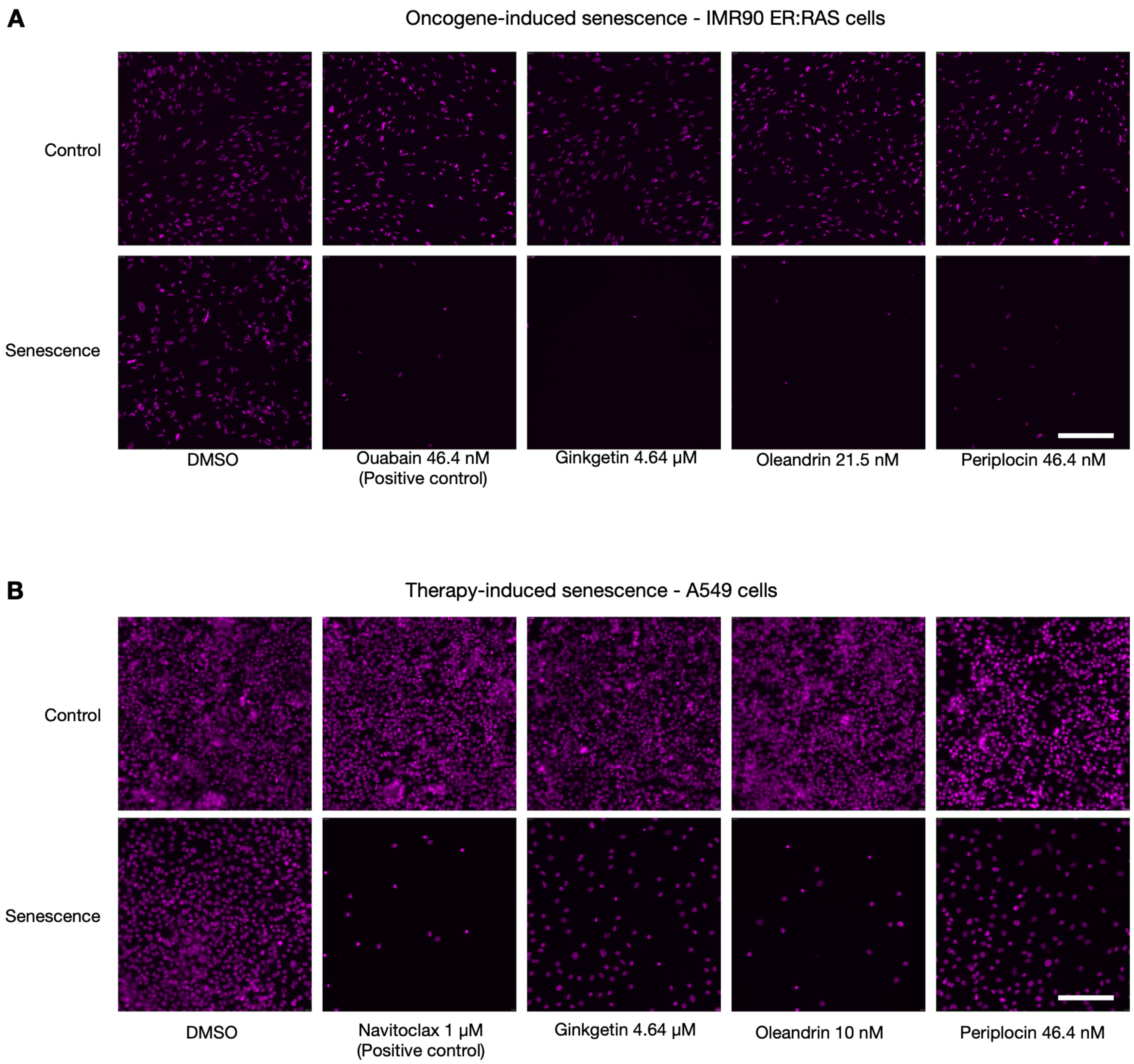


**Supplementary Figure 4. Representative images of nuclei for controls and newly found senolytics. (A)** 10× images of IMR90 ER:RAS cell nuclei of oncogene-induced senescence experiment. The positive control and the three found senolytics images are displayed at the minimal concentration where the senolytic effect was first observed (violet stain is DAPI channel). Scale bar is 100 μM. **(B)** 20× images of A549 cell nuclei of therapy-induced senescence experiment. The positive control and the three found senolytics images are displayed at the minimal concentration where the senolytic effect was first observed (violet stain is DAPI channel). Scale bar is 50 μM.

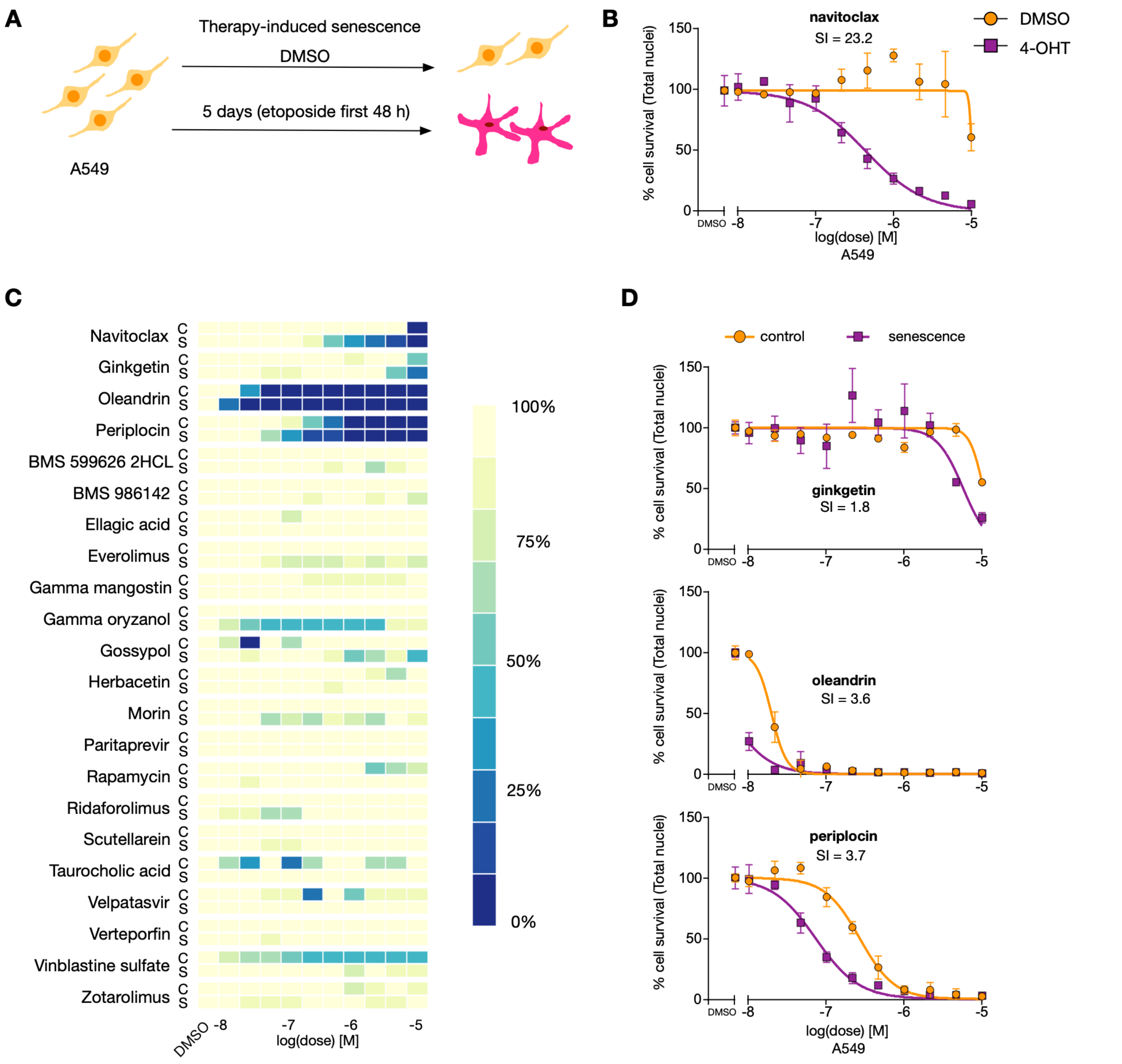


**Supplementary Figure 5. Experimental characterisation of compounds selected for screening in therapy-induced senescent cells. (A)** Experimental setup of therapy-induced senescence model with A549 cells. Senescence was induced by addition of etoposide at 100 μM for 48 hours, followed by three subsequent days of culturing in standard conditions. Control and senescent cells were plated on the last day of senescence induction. Top predicted compounds were added a day later, and 72 hours afterwards, the cells were fixed, and the nuclei stained and counted. **(B)** Dose-response curve of TIS positive experimental control, navitoclax. Data is normalised to DMSO. Mean ± s.d. are shown from n=3 experiments. **(C)** Results from experimental validation of controls and top 21 compounds predicted to have senolytic action with P>44%. **(D)** Dose-response curves of the three compounds out of 21 that displayed senolytic activity: ginkgetin, oleandrin, and periplocin. Data is normalised to DMSO. Mean ± s.d. are shown from n=3 experiments.

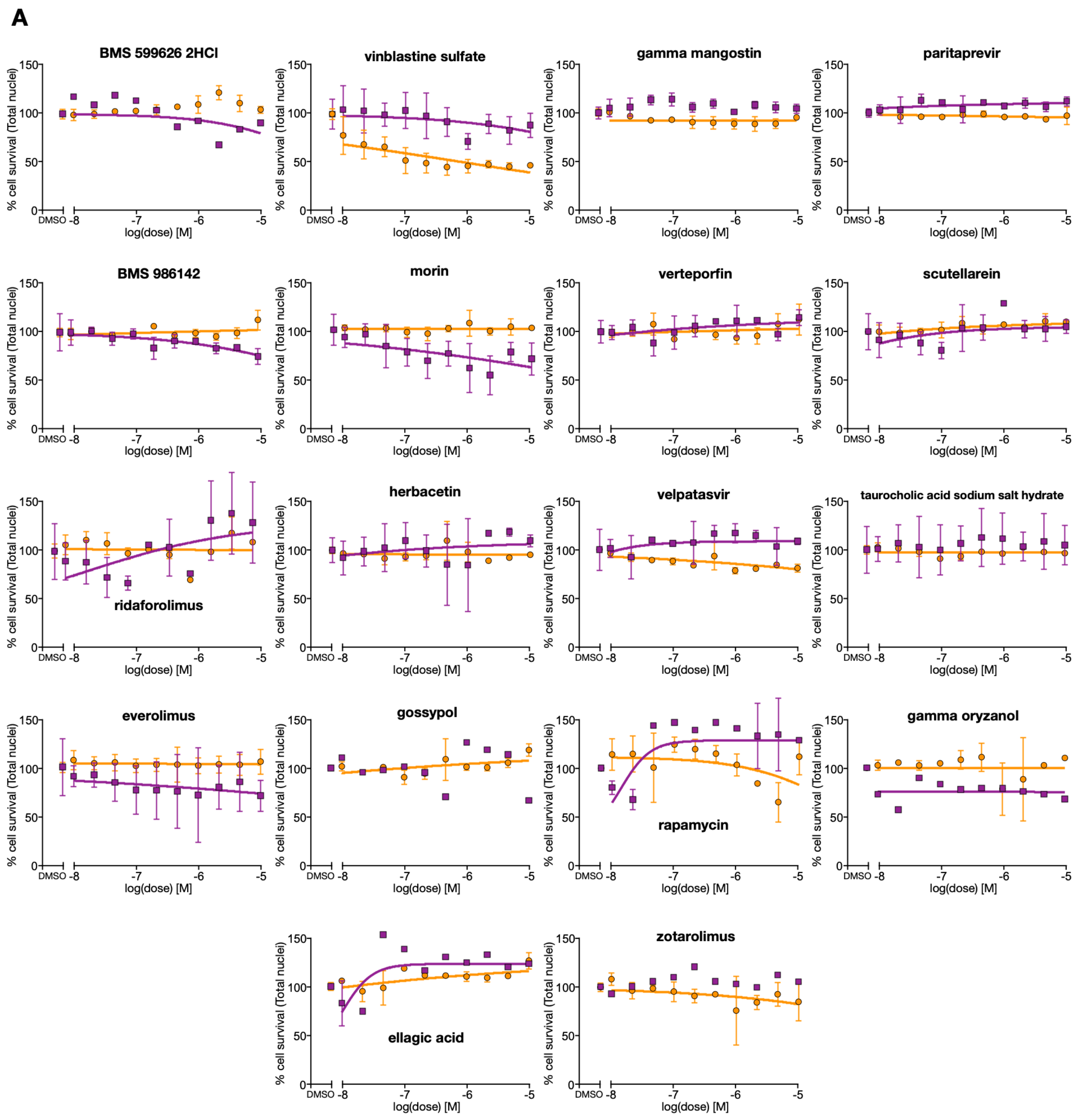


**Supplementary Figure 6. Experimental characterisation of false positives predicted by the machine learning model in therapy-induced senescent cells.** Dose-response curves of non-senolytic compounds predicted to have senolytic action by the XGBoost model with P>44%. Data is normalised to DMSO. Mean ± s.d. are shown from n=3 experiments.
